# Identification of Hyal2-expressing tumor-associated myeloid cells in cancer: implications for cancer-related inflammation through enhanced hyaluronan degradation

**DOI:** 10.1101/2020.09.14.296475

**Authors:** Paul R. Dominguez Gutierrez, Elizabeth P. Kwenda, William Donelan, Padraic O’Malley, Paul L. Crispen, Sergei Kusmartsev

## Abstract

Increased presence of myeloid derived suppressor cells (MDSCs) and tumor-associated macrophages (TAMs) in tumor tissue has been extensively reported. These cells represent a major constituent of tumor infiltrate and exhibit a distinct phenotype with immunosuppressive and tolerogenic functions. However, their role in the regulation of hyaluronan (HA) metabolism in the tumor microenvironment has not been established. Here we describe a novel function of tumor-associated myeloid cells related to the enhanced breakdown of extracellular HA in human bladder cancer tissue leading to accumulation of small HA fragments with MW <20 kDa. Increased fragmentation of extracellular HA and accumulation of low molecular weight HA (LMW-HA) in tumor tissue was associated with elevated production of multiple inflammatory cytokines, chemokines, and angiogenic factors. The fragmentation of HA by myeloid cells was mediated by the membrane-bound enzyme hyaluronidase 2 (Hyal2). The increased numbers of Hyal2^+^CD11b^+^ myeloid cells were detected in the tumor tissue as well as in the peripheral blood of bladder cancer patients. Co-expression of CD33 suggests that these cells belong to monocytic myeloid-derived suppressor cells. HA-degrading function of Hyal2-expressing MDSCs could be enhanced by exposure to tumor-conditioned medium, and IL-1β was identified as one of factors involved in the stimulation of Hyal2 activity. CD44-mediated signaling plays an important role in the regulation of HA-degrading activity of Hyal2-expressing myeloid cells, since engagement of CD44 receptor with specific monoclonal antibody triggered translocation of Hyal2 enzyme to the cellular surface and also stimulated secretion of IL-1β. Taken together, this work identifies the Hyal2-expressing tumor-associated myeloid cells, and links these cells to the accumulation of LMW-HA in the tumor microenvironment and cancer-related inflammation and angiogenesis.

## Introduction

The tumor stroma, which is comprised of cellular and extracellular components, plays a major role in tumor growth and progression. The extracellular matrix (ECM) of tumors includes proteoglycans and glycans such as hyaluronic acid, also called hyaluronan (HA). HA is a member of the glycosaminoglycan family of polysaccharides synthesized at the cell surface and is characterized by very high molecular weights (2×10^5^ to 10×10^6^ kDa) and extended lengths of 2-25 µm (**1**). Increased HA synthesis is associated with wound healing and tumor growth (**1-2**). Several cancer types including breast, prostate, brain, lung and bladder are highly enriched with HA (**2-4**). Within the tumor tissue, HA buildup is frequently associated with increased degradation of HA, leading to accumulation of low molecular weight HA (LMW-HA) fragments (**5, 6**). Several studies have demonstrated that LMW-HA displays unique biologic activities that are not shared by high molecular weight HA (HMW-HA) (**7-8**). HMW-HA is anti-oncogenic, anti-inflammatory and anti-angiogenic while, LMW-HA promotes inflammation and tumor angiogenesis by stimulating expression of cytokines, chemokines and growth factors in TLR2/TLR4 dependent manner (**9**). In addition, LMW-HA is a potent inducer of cPLA2 activity in macrophages that promotes release of arachidonic acid; a substrate for inflammation-associated lipid mediators PGE_2_ and leukotrienes **(9**).

It has been shown that progression of bladder cancer is associated with enhanced expression of hyaluronidase 2 (Hyal2) RNA in tumor tissue (**10, 11**). Hyal2, a member of hyaluronidase family, is a glycosylphosphatidylinositol-linked (GPI-linked) enzyme that is anchored to the plasma membrane and is involved in the degradation of extracellular HA (**12, 13**). Hyal2 cleaves high molecular weight HA into intermediate size 20 kDa fragments. In addition to increased Hyal2 expression, bladder cancer tissue is frequently infiltrated with inflammatory and immune cells (**14, 15**). Here we demonstrate that bladder tumor-associated myeloid cells express membrane-bound enzyme Hyal2. We also show that enhanced HA degradation in human bladder cancer is accompanied by the elevated production of inflammatory cytokines/chemokines and increased production of tumor angiogenic factors.

## Materials and Methods

### Human subjects

Freshly excised bladder tumor tissue and peripheral blood from 30 patients diagnosed with urothelial carcinoma of the bladder were collected during cystectomy or transurethral resection of bladder tumor (TURBT). Samples of normal bladder tissue were collected from patients undergoing cystectomy. All samples were obtained according to federal guidelines and as approved by the University of Florida institutional review board (IRB).

### Reagents and culture medium

The proteinase K and Benzonase were purchased from Sigma-Aldrich. Commercial polydisperse hyaluronan samples HA1M (MW 750-1000 kDa), 700K (500-749 kDa), 500K (301-450 kDa), 200K (151-300 kDa), 10K (10-20 kDa) and 5K (<10 kDa) were obtained from Lifecore Biomedical (Chaska, MN). Biotinylated hyaluronan-binding protein (HABP) wassupplied by Millipore-Sigma. Streptavidin conjugated with PE or FITC was purchased from Biolegend (San Diego, CA). Hyaluronan ELISA kit was purchased from R&D Systems. Hyaluronidase 2 polyclonal antibody conjugated with PE or Alexa-488 was obtained from Bioss Antibodies. All other antibodies used for immune fluorescence and flow cytometric analysis and human IL-1beta ELISA kit were acquired from Biolegend. Human was obtained from *In vitro* experiments were conducted using complete culture media consisting of RPMI 1640 medium supplemented with 20 mM HEPES, 200 U/ml penicillin, 50 μg/ml streptomycin (all from Hyclone) and 10% FBS from ATCC (Manassas, VA).

### Preparation of tissue slices from human normal and bladder cancer tissues

The precision-cut tissue slices, 2-4 mm in diameter and 200-300-micron thick, were produced from cancer and normal bladder tissues using a Compresstome vibratome VF-300-0Z. After cutting, tissue slices were placed into 24-well cell culture plates in complete RPMI-1640 medium supplemented with 10% FBS and antibiotics and cultured at 37° C in a humidified CO_2_ incubator. Cell viability of cultured tissue slices was tested using the Live/Dead cell viability kit purchased from Invitrogen.

### Isolation of CD11b myeloid cells from peripheral blood of cancer patients

PBMCs were isolated from bladder cancer patients by gradient density centrifugation using Lymphoprep (Accu-Prep, 1.077g/ml, Oslo, Norway). CD11b myeloid cells were purified from PBMCs by positive selection using the anti-CD11b microbeads and columns (Miltenyi Biotec). Briefly, cells were incubated with beads conjugated with anti-mouse CD11b and positively selected on LS columns. Viability of all recovered cells was 95%, as determined by trypan blue exclusion.

### Human cancer cell lines

The human T24 bladder cancer cell line was purchased from the ATCC (Manassas, VA). Tumor cells were maintained at 37°C in a 5% CO_2_ humidified atmosphere in complete culture medium.

### Preparation of tumor-conditioned medium

The source of tumor-conditioned medium (TCM) was bladder cancer tissue slice cultures or cultured T24 cell line. To prepare TCM, conditioned medium was collected 2-3 days after tissue or cell culture initiation, centrifuged, aliquoted and stored at -80°C.

### Cytokine and chemokine profiling

Human bladder cancer and normal tissue slices were cultured in a humidified CO_2_ incubator at 37°C. For profiling of cytokines in tissue conditioned media, cell-free culture supernatants were collected and stored at -80° C. Presence of 105 proteins in supernatants was evaluated using human cytokine and chemokine XL proteome array kit from R&D Systems (Minneapolis, MN).

### Analysis of HA produced by human bladder cancer

#### Visualization of tumor-produced HA

Cancer or normal bladder tissue slices were cultured for 7-14 days in 24-well cell culture plates in a humidified CO_2_ incubator at 37° C to allow for production of HA. At the end of incubation, tissue-produced HA was found settled at the bottom of the culture plate wells. To monitor and visualize accumulation of tissue-produced HA fragments on the plastic surface, the tissue slices and culture medium were removed at different time points. The empty wells were washed with warm PBS and fixed with 4% formaldehyde for 30 min. After fixation, plate wells were washed with PBS containing 2% FBS and incubated overnight with biotinylated HA-binding protein (3 μg/ml, Calbiochem-EMD Millipore) at 4° C (**16**). Next day, after washing the wells with PBS containing 2% FBS, streptavidin conjugated with fluorochrome was added to the wells and incubated for 30 min at 4°C. Plates were then washed with PBS and the bottoms of the wells were visualized using EVOS (Invitrogen) or Lionheart (Biotek Instruments) immunofluorescent imaging microscopes.

#### Evaluation of HA size

HA size analysis was determined using polyacrylamide gel electrophoresis as described previously (**17)**. Briefly, conditioned medium from cancer and normal bladder cancer tissue slices was centrifuged, aliquoted and stored at -80° C. To prepare samples for HA size analysis, thawed samples were digested with proteinase K to remove proteins, benzonase for depletion of nucleic acids (RNA, DNA) and ethanol to extract lipids was added. Samples along with HA standards were then subjected for polyacrylamide electrophoresis. The tissue-produced HA was visualized on the gel by staining with “Stains All” dye (Sigma-Aldrich).

### Immunofluorescent microscopy and flow cytometry

Immunofluorescent staining and flow cytometry analysis was performed as described previously (**18, 19**). Image analysis was done using Gen 5 Prime v 3.08 software (Biotek Instruments). Flow cytometry data and microscope pictures shown are representative of at least two separate determinations.

### Western blotting

Cells were lysed in M-PER^®^ mammalian protein extraction reagent (Thermo Scientific) containing protease and phosphatase inhibitors. Whole-cell lysates (30 µg/lane) were subjected to 10% SDS-PAGE, and blotted onto PVDF membranes. Membranes were blocked for 1 hour at room temperature with 5% dry skimmed milk in TBS (20 mM Tris-HCl, pH 7.6, 137 mM NaCl plus 0.1%, v/v, Tween 20) and probed with appropriate primary antibodies overnight at 4°C. Membranes were washed and incubated for 1 hour at room temperaturewith secondary Ab conjugated with HRP. Results were visualized by chemiluminescence detection using a SuperSignal West Pico substrate (Thermo Scientific). To confirm equal loading membranes were stripped using Restore™ Western blot stripping buffer (Thermo Scientific) and re-probed with antibody against β-Actin (Santa Cruz Biotechnology, Inc).

### Statistical analysis

The statistical significance between values was determined by the Student *t* test. All data was expressed as the mean ± SD. Probability values ≥ 0.05 were considered non-significant.

## Results

### Enhanced HA degradation in bladder cancer tissue results in accumulation of LMW-HA fragments

Normal bladder tissues produced predominantly long, structured linear pericellular HA (**Fig.1a**), whereas the bladder cancer samples generated highly fragmented tissue-associated HA (**Fig.1b**).

**Figure 1.**
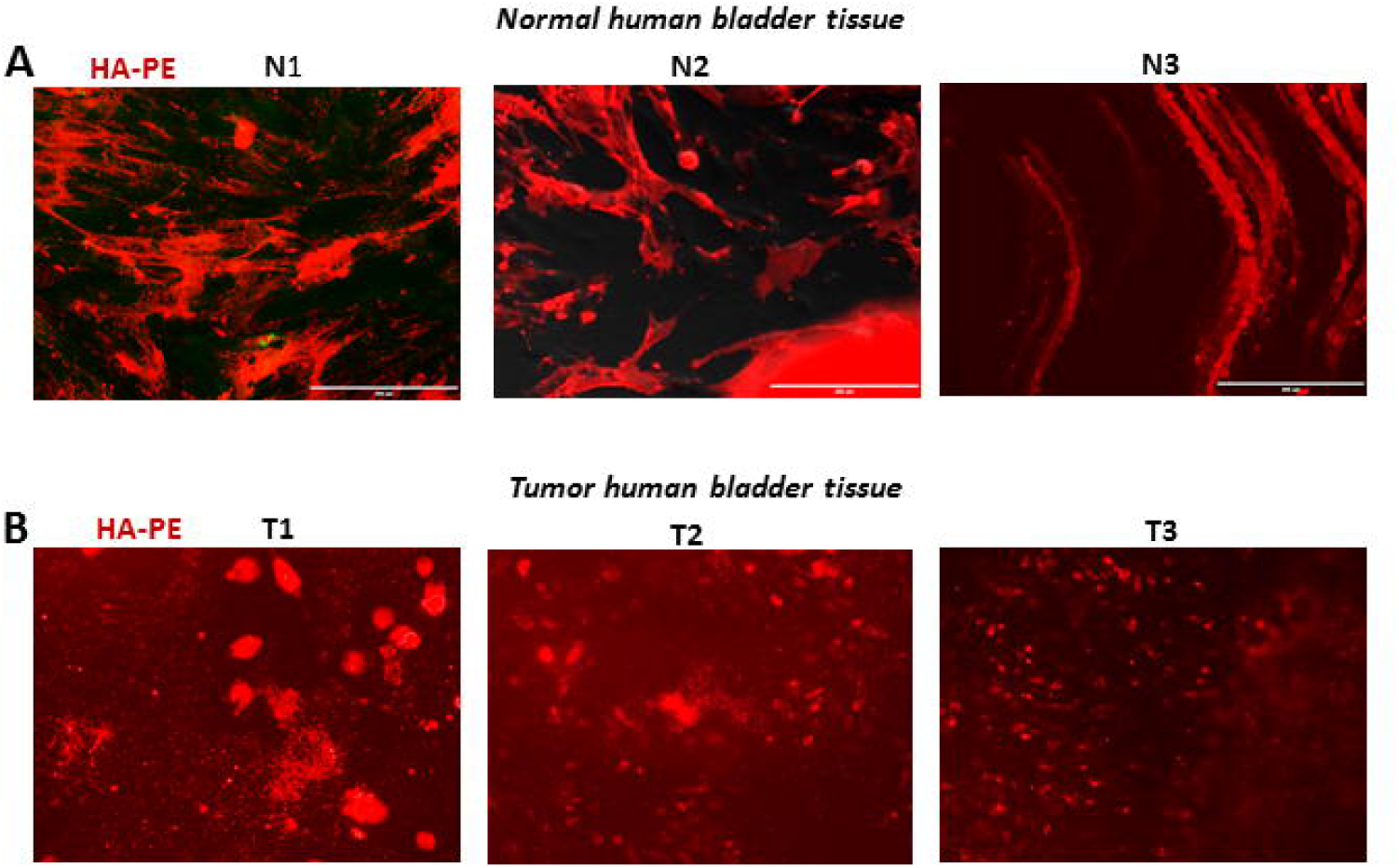
Visualization of HA produced by the tumor and normal human bladder tissue slices. Representative images showing tissue-produced HA in normal and tumor bladder tissues. Tissue slices prepared from normal (**A**) and tumor (**B**) human bladder tissue were cultured for 24 well plates for seven days. After removing culture medium and tissue slices, plates were fixed with 4% formaldehyde and stained for HA (red). Expression of HA was evaluated using IF microscopy.

The size of HA secreted by normal and bladder cancer tissues was further characterized by gel electrophoresis. Interestingly, tissues from normal bladder tissues (**Fig. 2a, left panel**) produced mostly HMW-HA (150 kDa and higher) with undetectable levels of LMW-HA. In contrast, the human bladder cancer tissues generated fragmented HA with a prevalence of LMW-HA with MW < 20 kDa (**Fig. 2a, right panel)** confirming our prior observation of increased HA fragmentation in bladder cancer tissues (**Fig.1b**).

**Figure 2.**
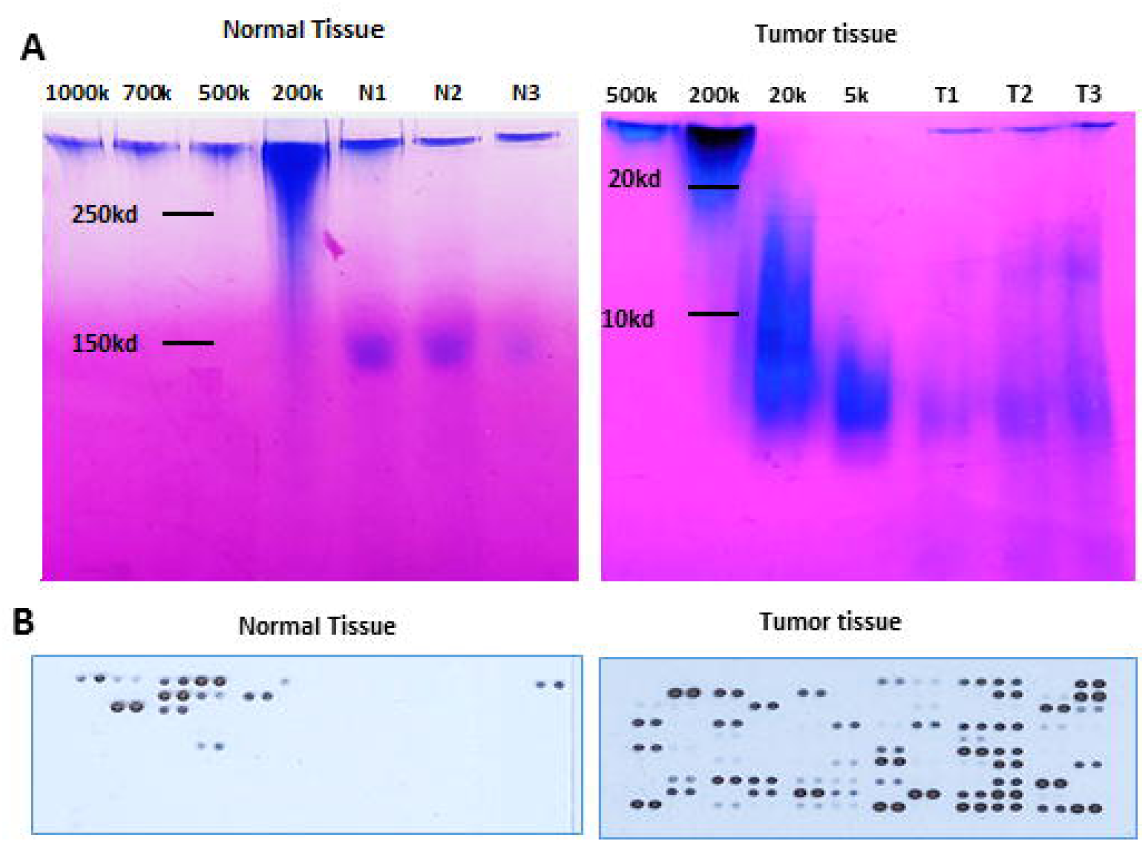
Enhanced HA degradation, accumulation of LMW-HA and elevated production of cytokines/chemokines in human bladder cancer tissue. Precision-cut tissue slices were prepared from freshly obtained normal and tumor human bladder tissue pieces and cultured in 24-well plates in full culture medium. Cell-free supernatants were collected on days 5-7, stored at 80° C until analysis of tumor-produced HA using polyacrylamide gel electrophoresis (**A**) and cytokines/chemokine antibody arrays (**B**).

### Enhanced HA degradation in tumor tissue is associated with elevated secretion of inflammatory, angiogenic and tumor-supporting factors

Accumulating evidence suggests that HA fragments with low molecular weight are able directly promote tumor progression by stimulating secretion of various factors that enhance tumor migration, invasion, inflammation, and angiogenesis. **(5, 6, 9**). To assess and compare the cytokine profile produced by normal and bladder cancer tissues, we analyzed the TCM collected from normal and bladder cancer tissue slices for the presence of 105 cytokines and chemokines using proteome multiplex assay. Our data revealed that normal bladder tissue (**Fig.2b**, left panel and **Supplementary Figs S1 and S2)** secreted detectable levels of nine proteins: adiponectin, ApoA1, chitinase3-like1, compliment factor D, C-reactive protein, endoglin, CXCL5, IL-8, and angiogenin. In TCM obtained from bladder cancer tissue, we detected 34 biologically active factors (**Fig.2b**, right panel and Supplementary Figures S1a, S1b). Among them, there were several angiogenic factors: VEGF, angiopoetin-2, angiogenin, and trombospondin-1; multiple chemokines and growth factors associated with inflammation and recruitment of different cell subsets to the tumor site such as CCL2, CCL7, CXCL1, CXCL5, CCL20, G-CSF, GM-CSF, lipocalin. The bladder cancer tissue also produced proteins involved in immune regulation (IL-6, IL-8, IL-11, IL-18Bpa, IL-24, IL-1R4, osteopontin) and tissue remodeling (MMP9, chitinase3-like1, dipeptyl-peptidase IV, IGFBP-2, uPAR, trombospondin). Proteins implicated in tumor cell invasion and migration, such as DKK1, cripto-1, HGF, were also detected. Taken together, this data suggests that accumulation of LMW-HA fragments within bladder cancer is accompanied by elevated production of multiple bioactive factors that are associated with tumor growth and progression.

### Detection of Hyal2-expressing tumor-associated myeloid cells in human bladder cancer

Increased levels of inflammatory chemokines and cytokines produced by the tumor facilitate cancer-associated inflammation and drive recruitment of myeloid cells to the tumor microenvironment where they become involved in bidirectional crosstalk with tumor cells (**20, 21**). As shown in **Fig. 3a**, bladder cancer tissue is infiltrated with myeloid cell subsets when compared to normal bladder tissue. Moreover, the tumor-infiltrating cells frequently associated with enhanced HA fragmentation (Fig. 3b and Supplementary Fig. S2). We hypothesized that the enhanced HA degradation within bladder cancer might be attributed to the presence and activity of tumor-recruited myeloid cells. To test this hypothesis, we co-stained HA produced by bladder cancer tissue slices with common myeloid cell marker CD11b. As shown in **Fig. 3d and Supplementary Fig. S3**, the CD11b^+^ cells localized in close proximity and some adhered to highly degraded HA fragments. Hyal2 is an enzyme responsible for HA degradation in the tumor microenvironment, whose increased expression in progressive bladder cancer was recently reported (**11**). Hence, we stained cancer tissue samples for the HA, Hyal2 and CD11b. Data presented in **Fig.3e and Supplementary Fig. S4** demonstrate the presence of Hyal2-expressing cells in tumor-bladder tissue slice cultures among both adherent and non-adherent cell fractions. Moreover, Hyal2 expression was observed in areas with highly fragmented HA and associated with a subset of tumor-infiltrating CD11b myeloid cells (**Fig.3f)** suggesting a possible contribution of these cells to the enhanced HA degradation in bladder cancer tissue.

**Figure 3.**
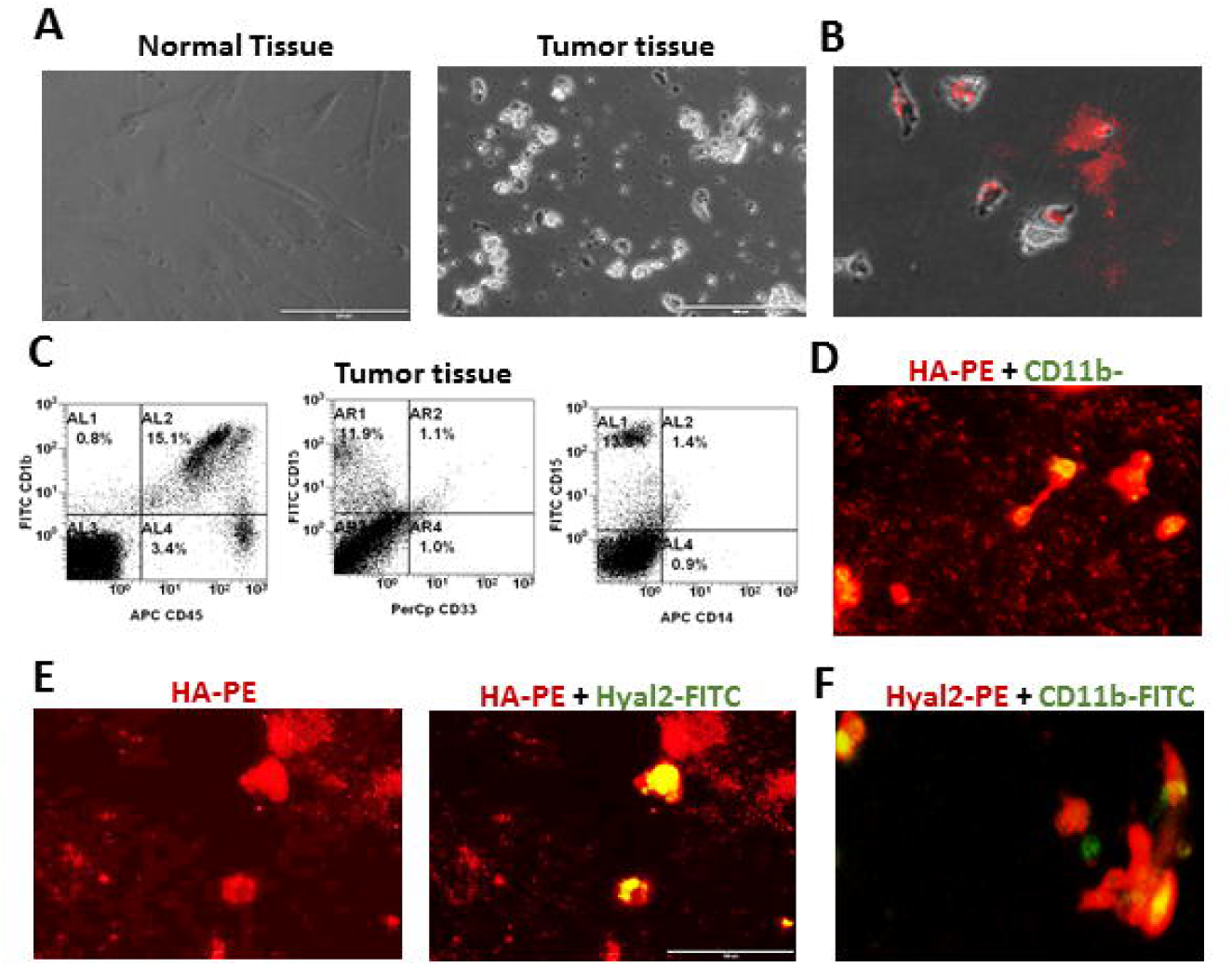
Degradation of HA in human tumor tissue is associated with the presence of tumor-infiltrating CD11b^+^Hyal2^+^ myeloid cells. **A**: Representative bright field images of normal bladder tissue (left panel) and bladder cancer tissue (right panel) from the same patient are shown. **B:** Co-localization of tumor-infiltrating cells and fragmented tumor-produced HA (red). Representative image is shown. **C**: Flow cytometric analysis of bladder cancer tissue. Tumor tissue from cancer patient was digested with collagenase cocktail to prepare single cell tumor suspension. The single cell suspension was co-stained with CD11b-FITC, CD45-APC, CD33-PerCp, CD15-FITC, CD14-APC antibodies and analyzed by flow cytometry. Representative images are shown. **D:** Co-localization of tumor-infiltrating CD11b myeloid cells and fragmented tumor-produced HA. Representative image of CD11b (green) and HA (red) in bladder cancer tissue is shown. **E:** Visualization of tumor-produced HA and tumor-associated Hyal2-expressing cells. The human cancer tissue slices were cultured for 5 days. Non-adherent cells were carefully removed, washed with PBS and fixed with 4% formaldehyde. To visualize the tumor-produced HA, biotinylated HA-binding protein and PE-labeled Streptavidin were subsequently added. Representative images of HA (red) and Hyal2 (green) are shown. **F**: Detection of tumor-infiltrating CD11b^+^Hyal2^+^ myeloid cells. Representative image of CD11b (green) and Hyal2 (red) in bladder cancer tissue is shown.

### Hyal2-expressing myeloid cells in peripheral blood of cancer patients

The peripheral blood of bladder cancer patients contain increased numbers of myeloid cells, including granulocytic and monocytic myeloid-derived suppressor cells (MDSCs) as compared to healthy individuals (**22**). To explore whether Hyal2-expressing myeloid cells could be also detected in peripheral blood, we isolated the CD11b^+^ cell populations from peripheral blood of bladder cancer patients and healthy donors, and stained with anti-Hyal2 Abs. Data presented in **Fig.4a and Supplementary Fig. S5** indicate that myeloid cells from blood of cancer patients contain significantly more Hyal2-positive cells as compared to healthy individuals. Additional analysis revealed that blood-derived Hyal2^+^ cells show mononuclear morphology (**Supplementary Fig. S6)**, also express marker of monocytic MDSCs CD33, as well as CCR2 receptor, M-CSF receptor (CD115) and GM-CSF receptor (CD16) (**Supplementary Fig. S7**).

**Figure 4.**
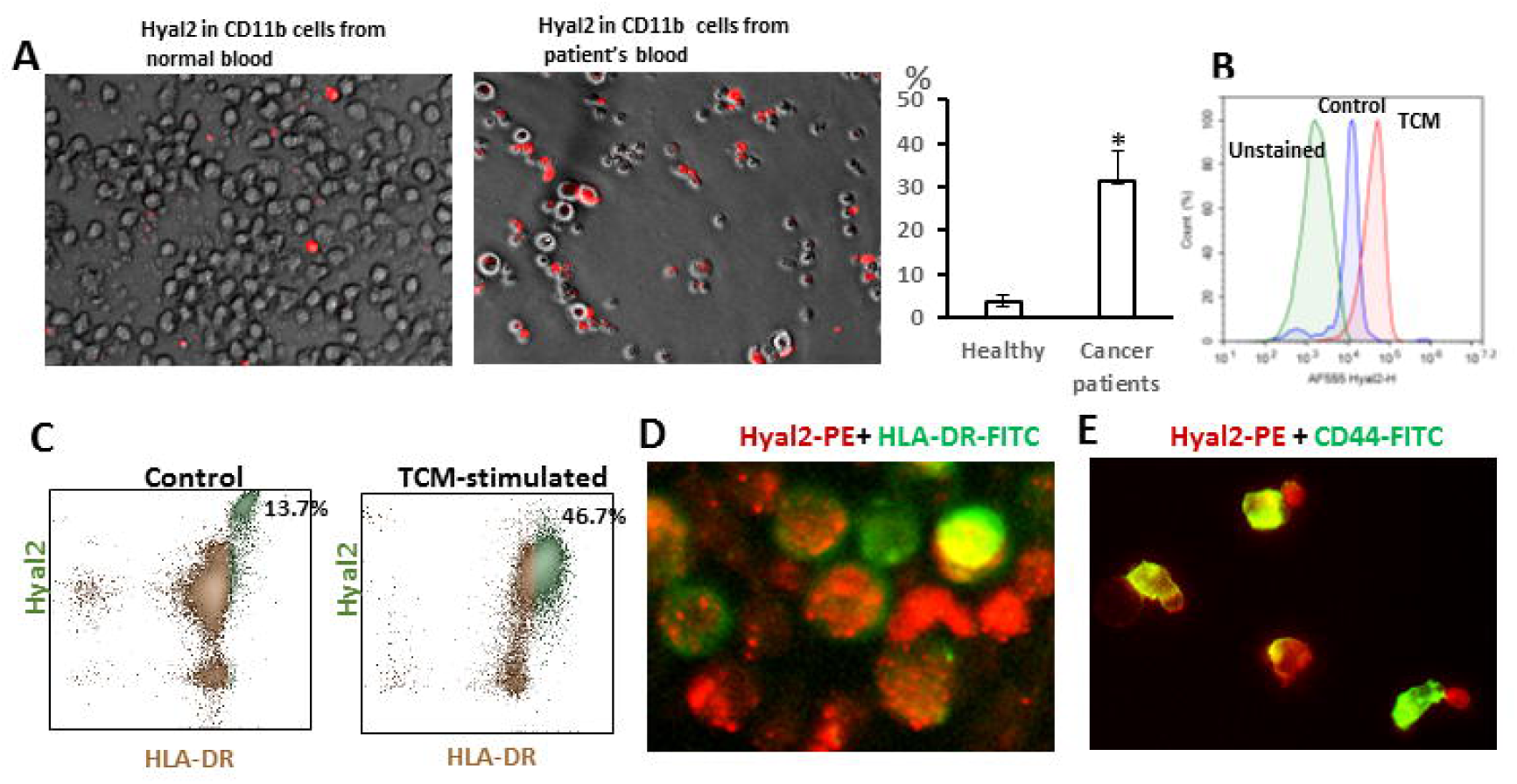
Identification of Hyal2-expressing myeloid cell subsets in peripheral blood from bladder cancer patients. **A:** Up-regulated expression of Hyal2 by peripheral blood-derived CD11b myeloid cells from cancer patients. CD11b myeloid cells were isolated from the peripheral blood of normal individuals or cancer patients using magnetic beads, stained with anti-Hyal2-PE antibodies and analyzed by IF microscopy. Percent of Hyal2^+^ cells was evaluated using immunofluorescent imaging microscope. Average means ± SD are shown; *, P<0.05 **B, C:** Analysis of Hyal2 expression in blood-derived myeloid cells using flow cytometry. CD11b myeloid cells were isolated from the peripheral blood of cancer patients and cultured in complete culture medium for 48 hours in the presence or absence of TCM. T24 tumor cell-derived culture supernatant was a source of TCM in these experiments. Collected cells washed with PBS and stained with anti-Hyal2 Abs (**B**), anti-Hyal2 and anti-HLA-DR abs (**C**). Expression of indicated markers was measured using flow cytometry. Representative images are shown. **D:** Analysis of Hyal2 localization in TCM-stimulated myeloid cells using IF microscopy. TCM-stimulated myeloid cells were prepared as indicated above and stained with anti-Hyal2-PE (red) and HLA-FITC (green) antibodies. Representative images are shown. **E:** Hyal2^+^ myeloid cells co-express CD44. CD11b myeloid cells were isolated from the peripheral blood of cancer patients using magnetic beads. TCM-stimulated myeloid cells were prepared as indicated above and stained with CD44-FITC and anti-Hyal2-PE. Representative image is shown.

We also found that exposure of myeloid cells to TCM further stimulated expression of Hyal2 **(Fig.4b)**, suggesting that expression of this enzyme could be up-regulated in myeloid cells upon recruitment to the tumor microenvironment with abundance of tumor-derived factors. Furthermore, a significant portion of Hyal2^+^ myeloid cells also co-expressed the antigen-presenting cell marker HLA-DR **(Fig.4c**): 13.7 % in non-stimulated vs 46.7% in TCM-stimulated CD11b myeloid cells. Interestingly, that in both TCM-treated (**Fig.4c**) and non-treated (data not shown) myeloid cells expression of Hyal2 was detected in dispersed granular form with both intracellular and cell surface localization (**Fig, 4d and Supplementary Fig. S7**). However, engagement of cells with antibodies against CD44 promoted significant changes in both cellular shape and Hyal2 localization (**Fig.4e**), supporting the idea that the CD44 receptor is involved in the regulation of Hyal2 function **(12, 13, 23, 24**).

### IL-1β stimulates HA-degrading activity of Hyal2^+^ myeloid cells

To examine whether TCM-stimulated CD11b^+^Hyal2^+^HLA-DR^+^ cells are functionally active, we tested their HA-degrading activity. To this end, we added the CD11b cells isolated from peripheral blood of cancer patients to the culture plates pre-coated with commercial HA (MW 200 kDa), and cultured cells in the presence of TCM for 10 days. Visualization of HA was executed using biotynalated hyaluronan-binding protein (HABP). As shown in **Fig. 5a (**right panel), TCM-stimulated myeloid cells were able to promote degradation of extracellular HA. Furthermore, by end of the culture (10 days), most of the TCM-stimulated myeloid cells acquired the shape of mature antigen-presenting cells and were able to internalize the fragmented HA (**Fig. 5b**). We next examined the potential cytokines/factors that could be involved in stimulation of HA-degrading activity mediated by Hyal2-expressing myeloid cells. Hyal2^+^ cells were isolated from peripheral blood of cancer patients and cultured in the presence or absence of following: human recombinant GM-CSF, M-CSF, IL-1β, osteopontin or TCM. HA-degrading activity of cultured Hyal2^+^ cells was evaluated by the visualization of degraded HA using IF microscopy (**Fig. 5c**), and quantification of small HA fragments detected on plastic surface in cell culture plates using imaging software (**Fig.5d**). Obtained data clearly demonstrate that only IL-1β among tested cytokines has been able to promote strong HA-degrading activity in Hyal2^+^ myeloid cells. Recently published study demonstrated that IL-1β can be produced by macrophages in CD44-dependent manner (**25**). Indeed, data presented in **Supplementary Fig. S**8 indicate that incubation of CD11b myeloid cells, isolated from peripheral blood of cancer patients, with anti-CD44 antibodies promoted the IL-1β secretion.

**Figure 5.**
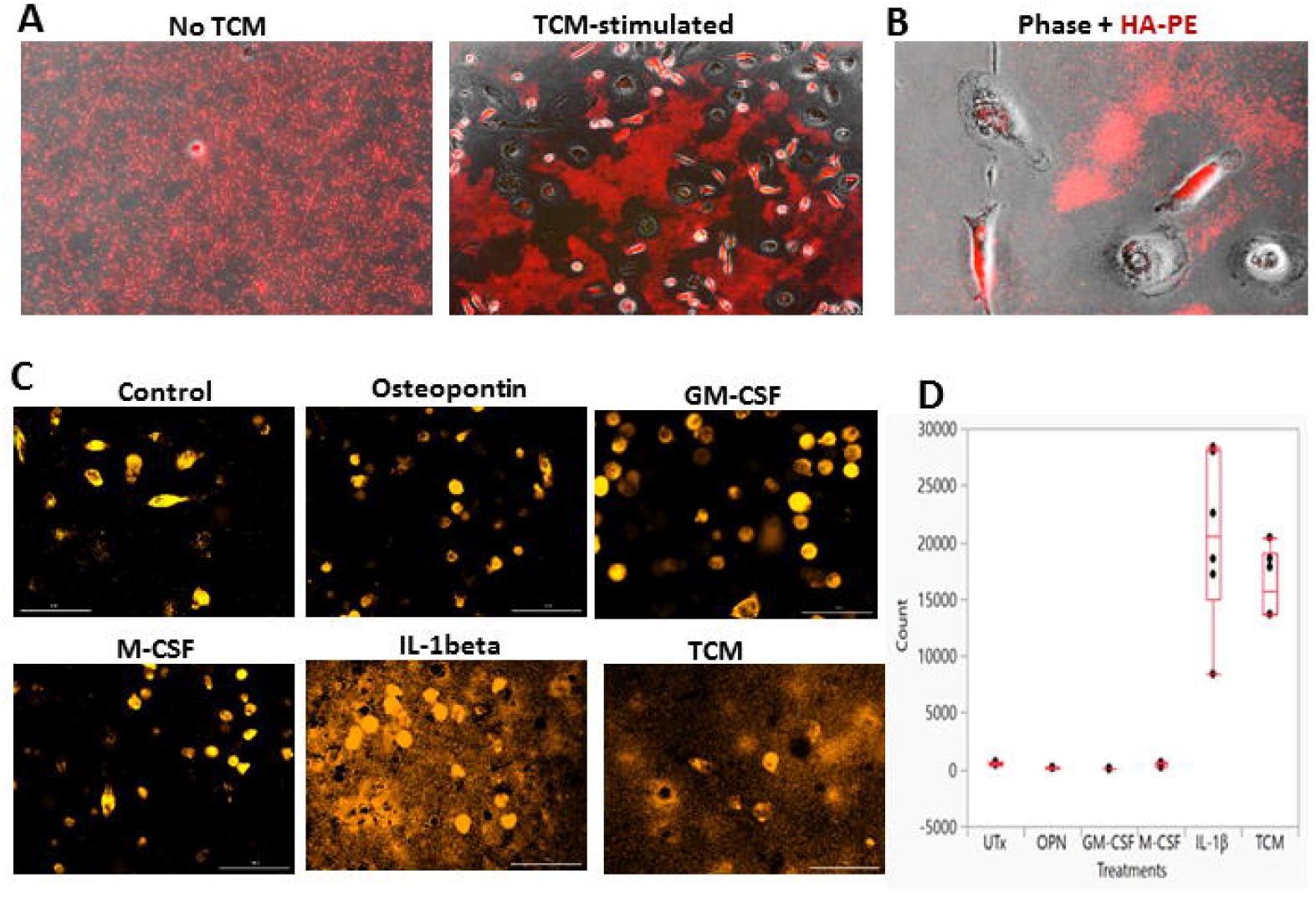
IL-1beta stimulates HA-degrading activity of Hyal2^+^ myeloid cells. **A:** Stimulation of peripheral blood-derived myeloid cells with TCM promotes degradation of extracellular HA. 24-well cell culture plates were pre-coated with sterile commercial HA (MW 200 kDa). CD11b myeloid cells isolated from peripheral blood of cancer patient were cultured in the absence (left panel) or in the presence of TCM (right panel). On day 10 cell cultures were fixed with 4% formaldehyde and stained for the HA (red). Representative images of control and TCM-treated myeloid cells cultured with HA are shown. **B:** Detection of intracellular HA in TCM-stimulated myeloid cells. CD11b myeloid cells isolated from peripheral blood of bladder cancer patient were added to the wells with pre-coated commercial HA. Cells were cultured in the presence of TCM for 10 days. Representative image is shown. **C**: IL-1beta stimulates HA-degrading activity of Hyal2^+^ myeloid cells. Hyal2^+^ were directly from peripheral blood of cancer patient and cultured in 24-well plate that was pre-coated with commercial HA (MW 200 kDa) in complete culture medium in presence of: bladder TCM, or GM-CSF (50 ng/ml), M-CSF (50 ng/ml), osteopontin (OPN, 50 ng/ml), IL-1beta (50ng/ml) or none (Utx). Supernatant from patient’s bladder tumor tissue culture was a source of TCM in these experiments. HA was visualized on Day 10 as described in section Materials and Methods using IF microscopy. **D**: Quantification of HA fragments in cytokine-treated Hyal2^+^ cells. Image analysis was done using Gen 5 Prime v 3.08 software (Biotek Instruments).

### Bone marrow as a source of Hyal2-expressing cells

Further analysis of myeloid cells obtained from peripheral blood of bladder cancer patients revealed that Hyal2-positive cells did not express the monocytic marker CD14 (**Fig 6a**). Long-term culture of Hyal2^+^ cells showed that these cells retained the antigen-presenting cell marker HLA-DR, but expression of Hyal2 becomes somewhat weaker and its localization was more intracellular overtime (**Fig.6b**). Importantly, the HLA-DR^+^CD11b^+^ cells and Hyal2^+^HLA-DR^+^ myeloid cells can be detected in bladder cancer tissue (**Fig. 6c and 6d, respectively**) in close proximity to areas that enriched for highly fragmented HA (**Fig.6d, right image**), suggesting the contribution of this myeloid cell subset to the process of HA degradation in the tumor microenvironment. Finally, taking into consideration that the primary location in the body where myelopoiesis takes place is bone marrow, we hypothesized that bone marrow may potentially serve as a source of Hyal2^+^ myeloid cells. Analysis of CD11b myeloid cells, isolated from normal human bone marrow, confirmed significance presence of Hyal2-expressing myeloid cells in bone marrow (**Fig. 6e**). Together, these data demonstrate that Hyal2-expressing myeloid cells can be detected in bladder cancer tissue, in close proximity to the HA degradation points.

**Figure 6.**
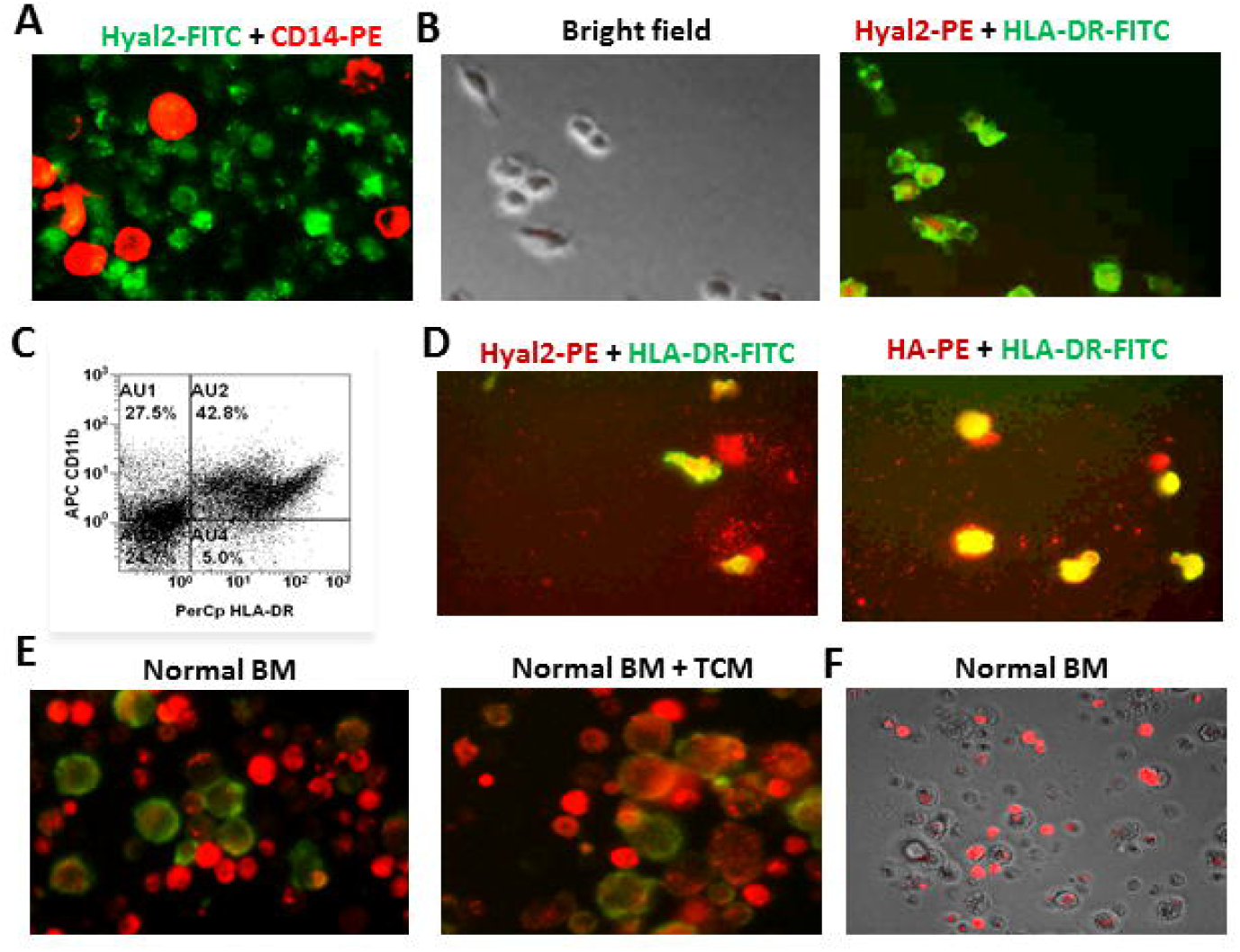
Bone marrow as a source of Hyal2-expressing cells. **A:** Hyal2 is not associated with CD14+ monocytes. CD11b myeloid cells were isolated from the peripheral blood of bladder cancer patients with magnetic beads and stained with antibodies against CD14 (red) and anti-Hyal2 (green). Representative image is shown. **B**: TCM drives development of Hyal2^+^HLA-DR^+^ cells using magnetic beads.. CD11b myeloid cells were isolated from the peripheral blood of bladder cancer patients and cultured in the presence of TCM in complete culture medium. On day 10, plates were washed with PBS, then fixed and stained with fluorochrome-conjugated antibodies against Hyal2 (red) and HLA-DR (green). Representative images are shown. **C:** Analysis of tumor-associated CD11b, HLA-DR-expressing cells using flow cytometry. Tumor tissue from bladder cancer patients were digested with collagenase cocktail to prepare single cell tumor suspension. The prepared cell suspension was co-stained with fluorochrome-conjugated antibodies against CD11b and HLA-DR and analyzed using flow cytometry. Representative image is shown. **D:** Detection of tumor-infiltrating Hyal2^+^HLA-DR^+^ cells. 24-well plates with human bladder cancer tissue slices were prepared for analysis of tumor-produced HA as described in section Materials and Methods. Representative images of HA-PE (red) adherent HLA-DR^+^ cells (green) are shown. **E:** Hyal2^+^ myeloid cells are enriched in bone marrow. CD11b myeloid cells were isolated from human normal bone marrow using magnetic beads. Non-stimulated (left panel) or TCM-stimulated (central and right panels) myeloid cells were prepared as indicated above and stained with anti-Hyal2-PE and HLA-DR-FITC antibodies. Representative images are shown.

## DISCUSSION

HA is a major component of the extracellular and pericellular matrix, which supports the normal tissue homeostasis. Deregulated HA metabolism has been observed in bladder, prostate, breast, brain, lung and other cancers. These cancers are characterized by elevated levels of HA and increased HA fragmentation in tumor tissue due to enhanced activity of hyaluronidases. Accumulating evidence suggests that HA fragments with LMW-HA directly stimulate secretion of various factors that promote tumor progression. However, the underlying mechanisms of increased HA degradation within the tumor microenvironment are not fully understood. Herein, we evaluated HA metabolism in bladder cancer samples obtained from patients undergoing surgical resection. Our analysis revealed that normal human bladder tissue produced HMW-HA, low levels of cytokines and contained negligible inflammatory and immune infiltrates. In contrast, bladder cancer tissues displayed strong HA degradation resulting in the accumulation of LMW-HA fragments and elevated production of chemokine, cytokines and angiogenic factors.

The molecular weight and size of HA fragments are important for the biological activity mediated by HA. HMW-HA has been shown to be anti-oncogenic (**7, 8**) and anti-inflammatory (**26**). In naked-mole rats, which display exceptional cancer-free longevity and a lifespan exceeding 30 years, extremely high molecular mass HA was found (**7**). This HMW-HA accumulates abundantly in naked mole-rat tissues due to decreased activity of HA-degrading enzymes and a unique sequence of HA producing enzyme hyaluronan synthase 2 (*HAS2)*. Once HMW-HA was removed by knocking down HAS2 or overexpressing the HA-degrading enzyme, Hyal2, naked mole-rat cells became susceptible to malignant transformation and readily formed tumors in mice. In stark contrast to HMW-HA, LMW-HA was shown to exert tumorigenic functions such as stimulation of cancer-related inflammation, tumor angiogenesis, and metastasis. Inflammation plays a critical role during different steps of tumor development and progression. Immune cells infiltrate tumors, creating an inflammatory microenvironment in which an extensive interaction -between immune cells and tumor cells takes place. LMW-HA fragments contribute to cancer-related inflammation by stimulating cytokine/chemokine production in a TLR2-and TLR4-dependent manner in both immune and tumor cells (**9, 27-29**). Furthermore, it was shown that degraded HA products with specific sizes of 4-25 disaccharides (MW ∼ 0,5-3,0 kDa) exert strong angiogenic activity (**29**). More recent studies have supported this finding (**31, 32**) by showing that CD44 and RHAMM-mediated signaling pathways are involved in HA-mediated angiogenesis. Additionally, several studies demonstrated that LMW-HA fragments have the pro-metastatic properties (**33, 34**).

In the present study, we demonstrate that myeloid cells in bladder cancer patients express the HA-degrading enzyme Hyal2. Membrane-bound enzyme Hyal2 exists as a glycosylphosphatidylinositol-linked (GPI-li2ked) protein exposed to the extracellular milieu (**12, 13, 35, 36**). It has been proposed that Hyal-1 and Hyal-2 are major mammalian hyaluronidases in somatic tissues, and that they act in concert to degrade HMW-HA to LMW-HA (**37-39**). 20-kDa intermediate size HA fragments generated by the Hyal-2 near the cell surface can be transported intracellularly and delivered to lysosomes, where Hyal-1 further degrades the 20-kDa to the LMW-HA fragments (< 5-10 kDa). Indeed, our observations support this model since majority of tumor-infiltrating or peripheral blood-derived Hyal2-expressing myeloid cells that exposed to the HA in the presence of activating factors (such as TCM or IL-1β) efficiently engulf HA fragments and show presence of the intracellular HA. On the other hand, significant amounts of LMW-HA fragments also were detected in extracellular space suggesting that both intracellular and extracellular small HA fragments could be involved in pathogenic LMW-HA-mediated signaling that stimulates enhanced production of inflammatory and angiogenic factors detected in the tumor tissue. In addition to cancer, Hyal2 has also been implicated in the pathogenesis of inflammatory joint diseases including rheumatoid arthritis and osteoarthritis. It was found that expression of Hyal2 in patients with arthritis is significantly up-regulated (**40**). Moreover, expression of Hyal2 in chondrocytes can be stimulated by IL-1β (**41**). Our data also support the direct role of IL-1β in stimulation of Hyal2-mediated HA degradation. IL-1beta is abundant in the tumor microenvironment and produced mostly by tumor-associated macrophages in response to the soluble CD44 (**43**). Recent study demonstrated that IL-1β supports both tumor progression and metastasis development (**43**). Orthotopic injection of murine mammary cells in IL-1beta knockout mice led to initial tumor growth but resulted in subsequent tumor regression and prevention of metastasis development. Moreover, treating mice first with anti-IL-1β Abs followed by anti-PD-1 antibodies completely abrogated tumor progression.

Some aggressive epithelial tumor cells show high expression of Hyal2 and able to degrade extracellular HA (**8**). However, Western Blotting analysis showed very weak expression of Hyal2 in human bladder cancer cell lines (Supplementary **Fig. S9**). Myeloid cells, including TAMs and MDSCs represent a major cellular component in tumor tissues that play a key role in tumor development and progression (reviewed in **20-22**). Recruitment of myeloid cells to the tumor microenvironment is a constant process fueled by the increased secretion of chemokines by both malignant as well as by stromal cells, which causes the mobilization of bone-marrow derived myeloid cell precursors from bone marrow and extravasation from the circulation into the tumor. Due to a tolerogenic cytokine milieu in the tumor microenvironment, recruited myeloid cells, differentiate into immunosuppressive TAMs and MDSCs. Tumor-recruited myeloid cells have been shown to exert supportive tumor-promoting effects via multiple pathways that stimulate local immune suppression/tolerance, tumor angiogenesis, tissue remodeling and cancer inflammation. However, their role in the degradation of extracellular HA and, particularly, of tumor-associated HA has not been recognized yet. We identified the myeloid cell subset within bladder cancer tissue that expresses the HA degrading Hyal2 suggesting involvement of these cells in the enhanced fragmentation of extracellular HA observed in tumor tissue. Increased presence of Hyal2-expressing myeloid cells was also detected in the peripheral blood of bladder cancer patients. HA-degrading function of Hyal2-expressing myeloid cell could be enhanced by exposure to the TCM, and IL-1β was identified as one of factors stimulating the Hyal2 activity. CD44-mediated signaling plays an important role in the regulation of HA-degrading activity of myeloid cells, since stimulation of CD44 receptor with specific monoclonal antibody triggered secretion of IL-1β and translocation of Hyal2 to the cellular surface. Collectively, this work identifies the Hyal2-expressing myeloid cells and links these cells to the accumulation of LMW-HA in tumor microenvironment.

## Authors’ Contributions

Conception and Design: S.K.

Acquisition of data (acquired and managed patients, provided facilities, provided animals): P.R.D.G, P.C., P.O.M., E.K, W.D.

Writing, review, editing of the manuscript: S.K, P.R.D.G, P.C., P.O.M., E.K, W.D. Administrative, technical or material support: S.K., P.C., P.R.D.G.

Study supervision: S.K.

## ACKNOWLEDGEMENTS

This work supported by grant #8JK05 from J&E King Biomedical Research Program and Fund 1923 to S.K.

## References

1. Girish, K.S., and K. Kemparaju. 2007. The magic glue hyaluronan and its eraser hyaluronidase: a biological overview. Life Sciences. 80:1921–1943

2. Toole, B.P. 2004. Hyaluronan: from extracellular glue to pericellular cue. Nat. Rev. Cancer. 4:528–539.

3. Sironen, R., M. Tammi, R. Tammi, P.K. Auvinen, M. Anttila, and V.M. Kosma.Hyaluronan in human malignancies. Exp. Cell Res, 2011, 317: 393–91.

4. Simpson M.A and V.B. Lokeshwar. Hyaluronan and hyaluronidase in genitourinary tumors. Front. Bioscience, 13, 2008, 5664–5680.

5. Schmaus A, Bauer J and J. P. Sleeman. Sugars in the microenvironment: the sticky problem of HA turnover in tumors. Cancer Metastasis Rev, 2014, 33:1059–1079

6. Turley E.A., Wood D.K, and McCarthy J.B. Carcinoma cell hyaluronan as a “portable” cancerized pro-metastatic microenvironment. Cancer Res, 2016, 76(9): 2507–2512.

7. Tian X et al. High-molecular-mass hyaluronan mediates the cancer resistance of the naked mole rat. Nature, 2013, (499): 346–349.

8. Ooki T, Murata-Kamiya N, Takahashi-Kanemitsu A., Wu W and Hatakeyama M. High-molecular-weight hyaluronan is a hippo pathway ligand directing cell density-dependent growth inhibition via PAR1b. Dev. Cell, 2019, 49(4):590–604.

9. Sokolowska, M., L. Y. Chen, M. Eberlein, A. Martinez-Anton, Y. Liu, S. Alsaaty, H.Y. Qi, C. Logun, M. Horton, and J.H. Shelhamer. 2014. Low molecular weight hyaluronan activates cytosolic phospholipaseA2α and eicosanoid production in monocytes and macrophages. J. Biol. Chem. 289:4470–4488.

10. Kramer M.W et al. HYAL-1 hyaluronidase: A potential prognostic indicator for progression to muscle invasion and recurrence in bladder cancer. Eur. Urol., 2010, 57:8 6 –9 4.

11. Van Der Heijden AG et al. A five-gene expression signature to predict progression in T1G3 bladder cancer. Eur J Cancer, 2016, 64:127–36.

12. Stern R. Hyaluronidases in cancer biology. Semin Cancer Biol, 2008, 18: 275–280.

13. Harada H and Takahashi M. CD44-dependent intracellular and extracellular catabolism of hyaluronic acid by hyaluronidase-1 and -2. J. Biol. Chem., 2007, 282 (8): 5597–5607.

14. Eruslanov E, McCullers M, Daurkin I, Algood C, Dahm P, Rosser CJ, Vieweg J, Gilbert SM Kusmartsev S. Circulating and tumor-infiltrating myeloid cell subsets in patients with bladder cancer. International Journal of Cancer, 2012, 130(5):1109–19.

15. Puttman K. et al., The role of myeloid derived suppressor cells in urothelial carcinoma Immunotherapy. bladder cancer, 2019.5 (2):103–11

16. Petrey, A.C., and C.A. de la Motte. 2016. Thrombin cleavage of inter-α-inhibitor heavy chain 1 regulates leukocyte binding to an inflammatory hyaluronan matrix. J Biol. Chem. 291:24324–24334.

17. Min H, Cowman MK. Combined Alcian Blue and silver staining of glycosaminoglycans in polyacrylamide gels: application to electrophoretic analysis of molecular weight distribution. Anal. Biochem. 1986;155: 275–285.

18. Daurkin I, Eruslanov E, Stoffs T, Perrin GQ, C. Algood, Gilbert SM, Rosser CJ, Su L.M, Vieweg J, Kusmartsev S. Tumor-associated macrophages mediate immune suppression in kidney cancer microenvironment by activating 15-lipoxygenase pathway. Cancer Research, 2011, 71(20):6400–9.

19. Prima, V., L. Kaliberova, S. Kaliberov, D. Curiel D, and S. Kusmartsev. COX2-mPGES1-PGE2 pathway regulates PD-L1 expression in tumor-associated macrophages and myeloid-derived suppressor cells. PNAS, 2017, (114):1117–1122.

20. Biswas, S.K., and A. Mantovani. Macrophage plasticity and interaction with lymphocyte subsets: cancer as a paradigm. Nat. Immunol. 2010, 11:889–896.

21. Gabrilovich, D.I, S. Ostrand-Rosenberg, and V. Bronte. Coordinated regulation of myeloid cells by tumours. Nat. Rev. Immunol., 2012, 12:253–268.

22. Crispen PL and Kusmartsev S. Mechanisms of immune evasion in bladder cancer. Cancer Immunology and Immunotherapy, 2019, doi: 10.1007/s00262-019-02443-4.

23. Bourguignon LY, Singleton PA, Diedrich F, Stern R, Gilad E. CD44 interaction with Na+-H+ exchanger (NHE1) creates acidic microenvironments leading to hyaluronidase-2 and cathepsin B activation and breast tumor cell invasion. J Biol Chem. 2004; 279:26991–27007.

24. Duterte C, Mertens-Strijthagen J, Tammi M, Flamion B. Two novel functions of hyaluronidase-2 (Hyal2) are formation of the glycocalyx and control of CD44-ERM interactions. J Biol Chem. 2009; 284:33495–33508.

25. Jang J.H et al. Breast Cancer Cell–Derived Soluble CD44 Promotes Tumor Progression by Triggering Macrophage IL1b Production. Cancer Res, 2020, 80:1342–1956.

26. Jiang, D., Liang, J., & Noble, P. W. (2011). Hyaluronan as an immune regulator in human diseases. Physiological Reviews, 91(1), 221–264.

27. Cnanmee T, Ontong P, Itano N. Hyaluronan: A modulator of the tumor microenvironment. Cancer Lett, 2016 375(1):20–30.

28. Jiang, D., Liang, J., Fan, J., Yu, S., Chen, S., Luo, Y. et al. (2005). Regulation of lung injury and repair by Toll-like receptors and hyaluronan. Nat Med, 11(11), 1173–1179.

29. Voelcker, V., Gebhardt, C., Averbeck, M., Saalbach, A. Wolf, et al. Hyaluronan fragments induce cytokine and metalloprotease upregulation in human melanoma cells in part by signalling via TLR4. Exp. Dermatol., 2008, 17(2), 100–107.

30. West DC, Hampson IN, Arnold F and Kumar S. Science, 1985, 228 (4705):1324–6.

31. Matou-Nasri, S., Gaffney, J., Kumar, S., & Slevin, M. Oligosaccharides of hyaluronan induce angiogenesis through distinct CD44 and RHAMM-mediated signalling pathways involving Cdc2. International Journal of Oncology, 2009, 35(4), 761–773.

32. Gao, F., Liu, Y., He, Y., Yang, C., Wang, Y., Shi, X., & Wei, G. Hyaluronan oligosaccharides promote excisional wound healing through enhanced angiogenesis. Matrix Biology: Journalof the International Society for Matrix Biology, 2010, 29(2): 107–116.

33. Itano, N., Sawai, T., Miyaishi, O., & Kimata, K. Relationship between hyaluronan production and metastatic potential of mouse mammary carcinoma cells. Cancer Res, 1999, 59(10): 2499–2504.

34. Schmaus, A., Klusmeier, S., Rothley, M., Dimmler, A., Sipos, B., Faller, G. et al. Accumulation of small hyaluronan oligosaccharides in tumour interstitial fluid correlates with lymphatic invasion and lymph node metastasis. Br J of Cancer, 2014, 2014.332

35. Rai SK, Duh FM, Vigdorovich V, Danilkovitch-Miagkova A, Lerman MI, Miller AD. Candidate tumor suppressor HYAL2 is a glycosylphosphatidylinositol (GPI)-anchored cell-surface receptor for jaagsiekte sheep retrovirus, the envelope protein of which mediates oncogenic transformation. Proc Natl Acad Sci USA. 2001; 98:4443–4448.

36. Andre B, Duterme C, Van Moer K, Mertens-Strijthagen J, Jadot M, Flamion B. Hyal2 is a glycosylphosphatidylinositol-anchored, lipid raft-associated hyaluronidase. Biochem Biophys Res Commun. 2011; 411:175–179.

37. Stern R. Devising a pathway for hyaluronan catabolism: are we there yet. Glycobiology. 2003, 105–115.

38. Czoka A.b et al. The six hyaluronidase-like genes in the human and mouse genomes. Matrix Biol, 2001 20:499–508.

39. Stern R. Hyaluronan metabolism: a major paradox in cancer biology. Pathol Biol (Paris), 2005, 53(7):372–82.

40. Yoshida M et al, Expression analysis of three isoforms of hyaluronan synthase and hyaluronidase in the synovium of knees in osteoarthritis and rheumatoid arthritis by quantitative real-time reverse transcriptase polymerase chain reaction. Arthritis Res. Ther., 2004, 6:514–20.

41. Tanimoto k et al. Modulation of hyaluronan fragmentation by interleukin-1beta in synovial membrane cells, Ann Biomed. Eng. 2010, 38:1618–25

42. Jang JH, Kim DH, Lim JM, Lee JW, Jeong SJ, Kim KP and Surh YJ. Breast cancer cell-derived soluble CD44 promotes tumor progression by triggering macrophage IL-1β production. Cancer Res, 2020, 80 (6):1342–56.

43. Kaplanov I, Carmi Y, Kornetsky R, Shemesh A, Shurin GV, Shurin MR, Dinarello CA, Voronov E, Apte RN. Blocking IL-1β reverses the immunosuppression in mouse breast cancer and synergizes with anti-PD-1 for tumor abrogation. Proc Natl Acad Sci USA., 2019, 116(4):1361–1369

