## Supplemental Figures for "Identification of Hyal2-expressing tumor-associated myeloid cells in cancer: implications for cancer-related inflammation through enhanced hyaluronan degradation"

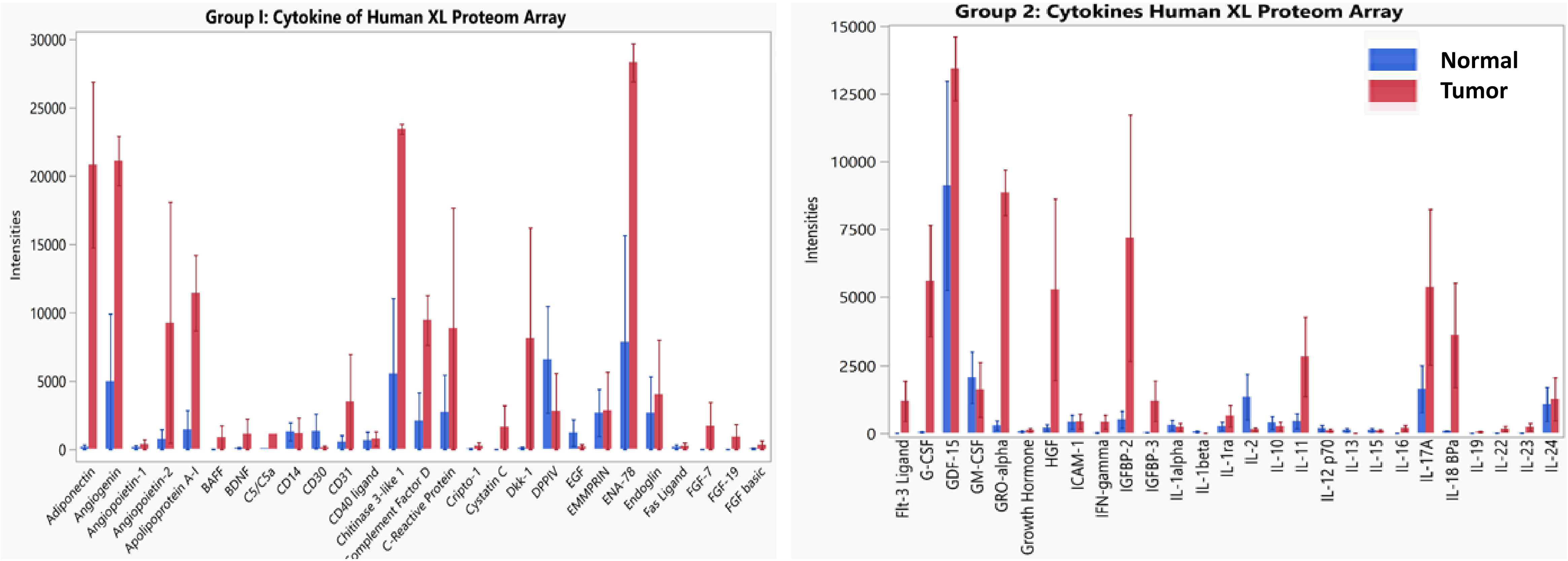


**Figure S1.** Precision-cut tissue slices were prepared from freshly obtained normal and tumor human bladder tissue pieces and cultured in 24-well plates in full culture medium. Cell-free supernatants were collected on day 5, stored at $80^0$ C until analysis of tumor-produced HA using cytokines/chemokine antibody arrays (**A**, **B**) from R&D Systems. Combined data from 3 normal and 3 tumor bladder tissue samples are shown. Quantification of the arrays was done with Quick Spot (Western Union Software) . Collected data were analyzed and graphed including the heatmap using JMP Pro 15.1.0 (SAS). The error bars show standard error for each analyte in triplicates.

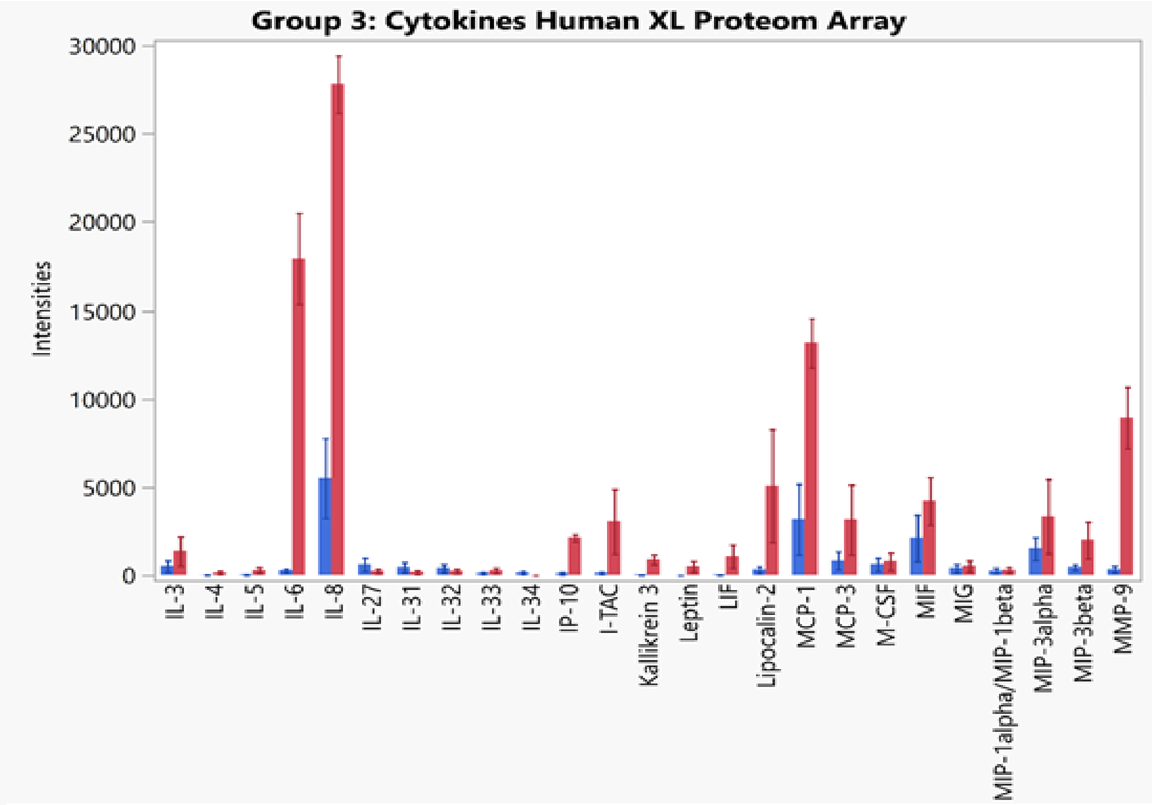


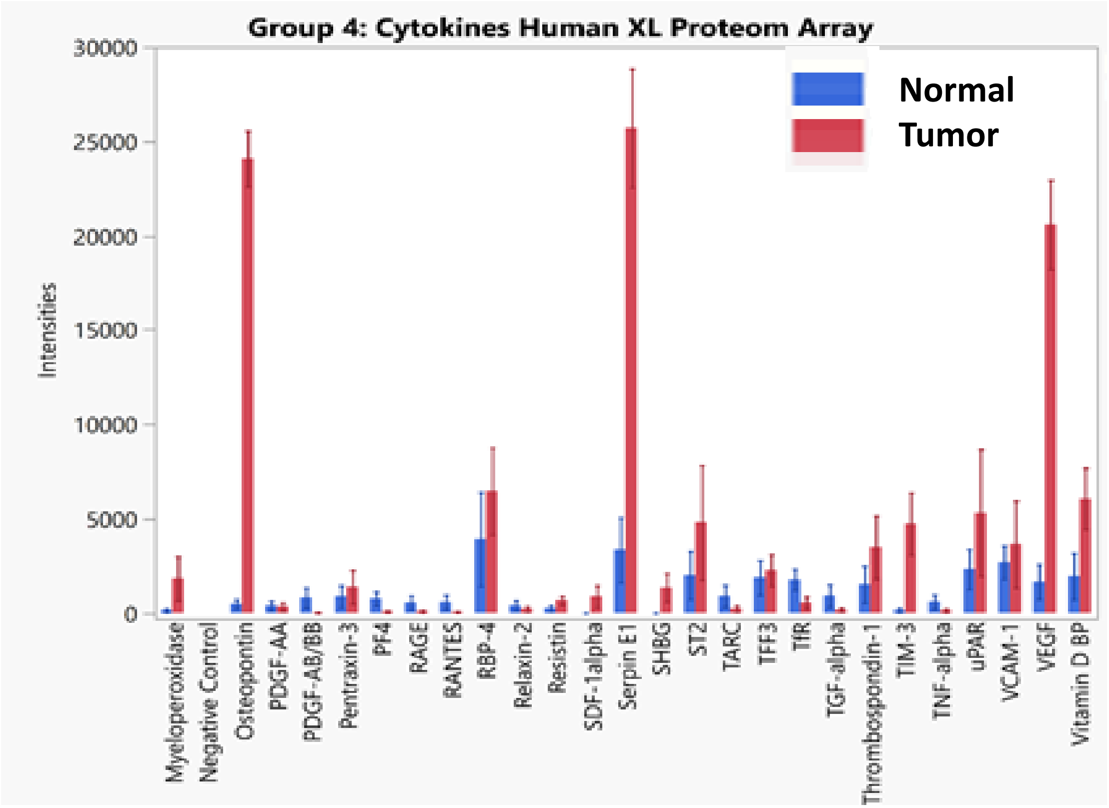


**Figure S1** (continuation).

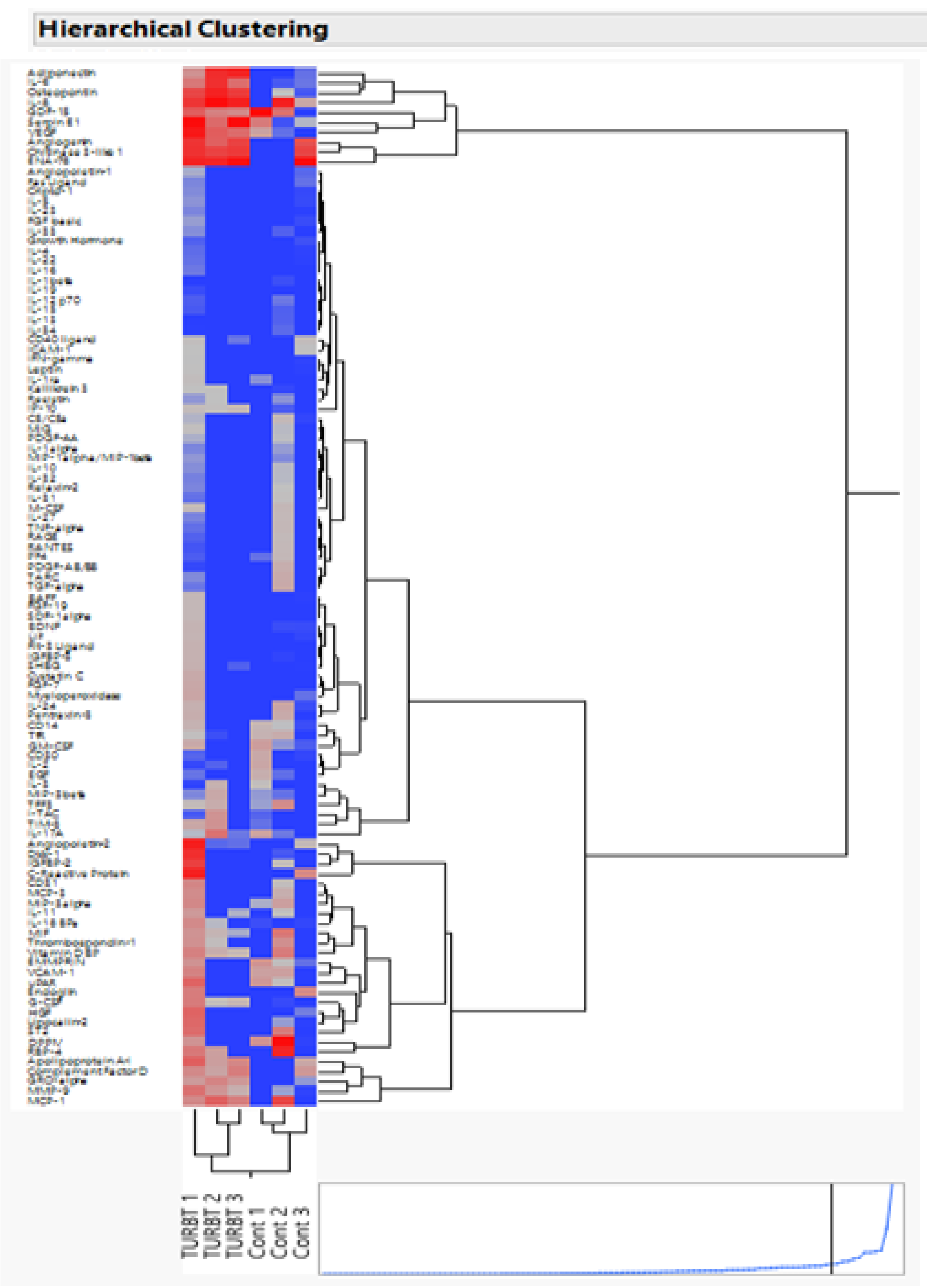


**Figure S2.** Hierarchical Clustering map/ Heatmap for factors produced by tumor (red) and normal (blue) bladder tissues.

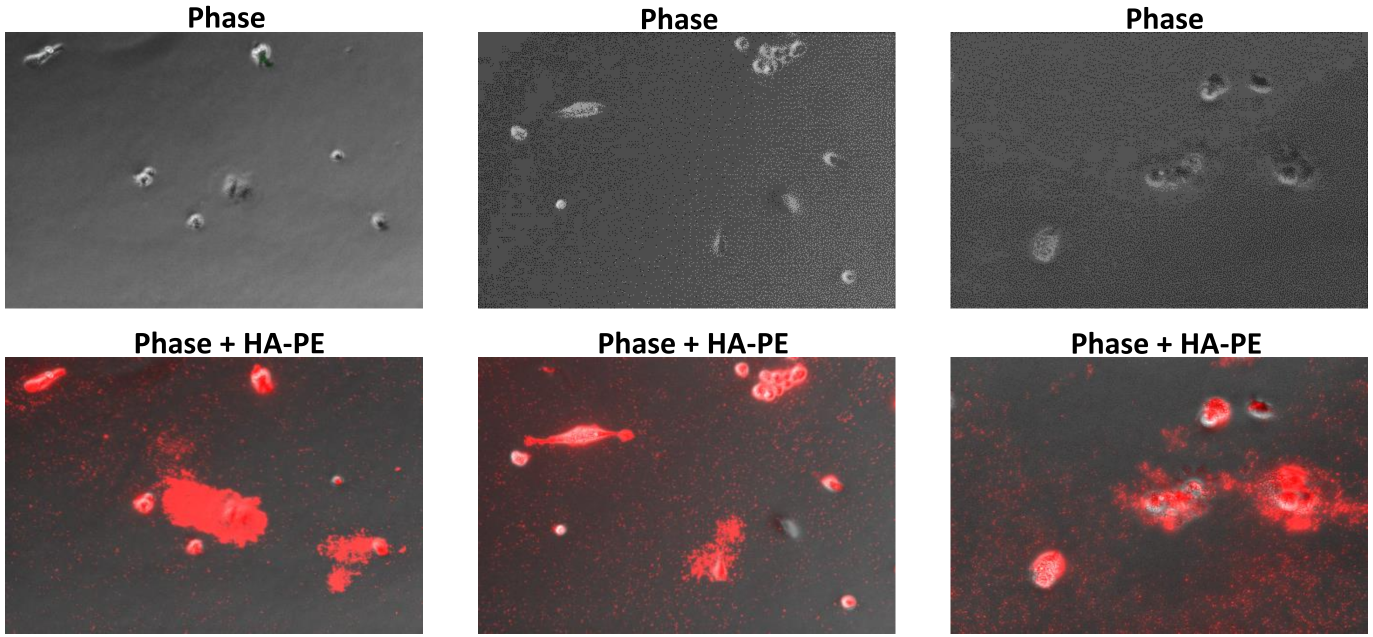


**Figure S3.** Co-localization of tumor-infiltrating cells and fragmented tumor-produced HA. Representative images of HA (red) and infiltrating cells (phase) in bladder tumor-tissue slice cultures are shown.

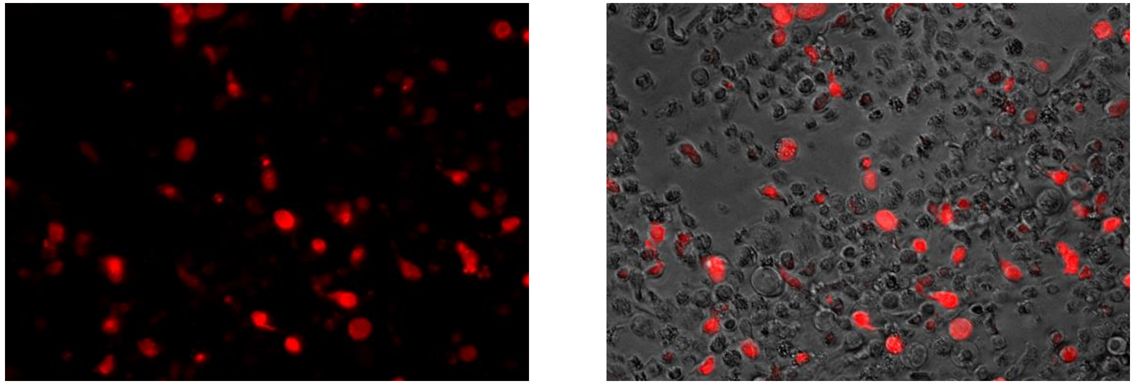

**Figure S4**. Detection of Hyal2-expressing cells in non-adherent fraction of cells obtained from bladder tumor tissue slice culture. Representative images of Hyal2 expression (red) in bladder cancer tissue is shown.

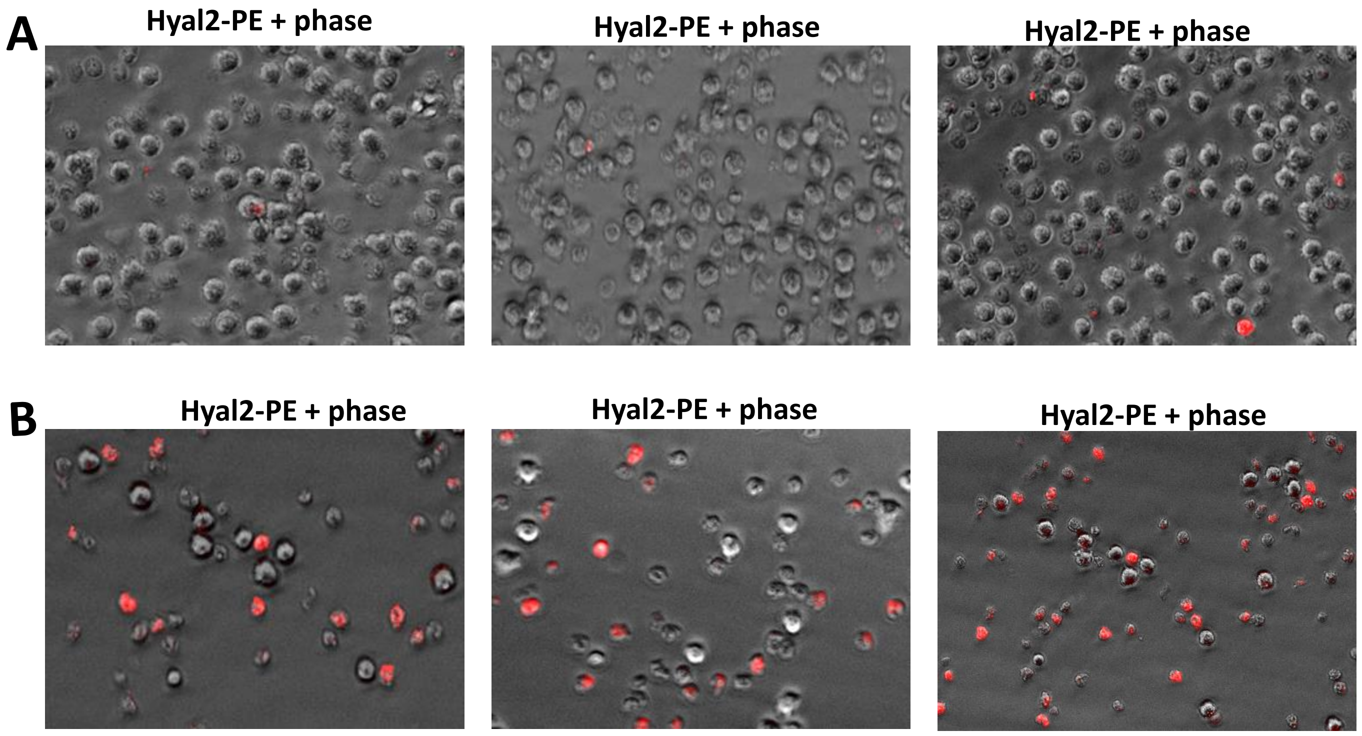


**Figure S5**. Hyal2 expression in CD11b cells isolated from blood of healthy donors (**A**) and bladder cancer patients (**B**). PBMCs were enriched from whole blood of 3 healthy individuals and 3 bladder cancer patients. CD11b myeloid cells were isolated from PBMCs by positive selection using magnetic beads and stained with anti-Hyal2-PE antibodies.

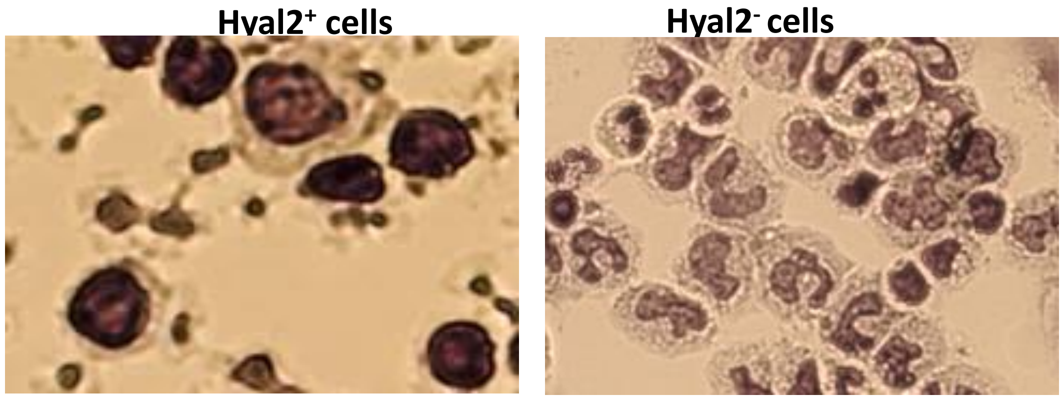


**Figure S6.** Hyal2$^+$ and Hyal2$^-$ CD11b$^+$ myeloid cells were sorted from peripheral blood of bladder cancer patients using magnetic beads. Cytospins with purified cells were prepared using Shandon cytospin centrifuge, fixed and stained with haematoxylin/eosin. Representative images of Hyal2$^+$ and Hyal2$^-$ CD11b$^+$ myeloid cells are shown.

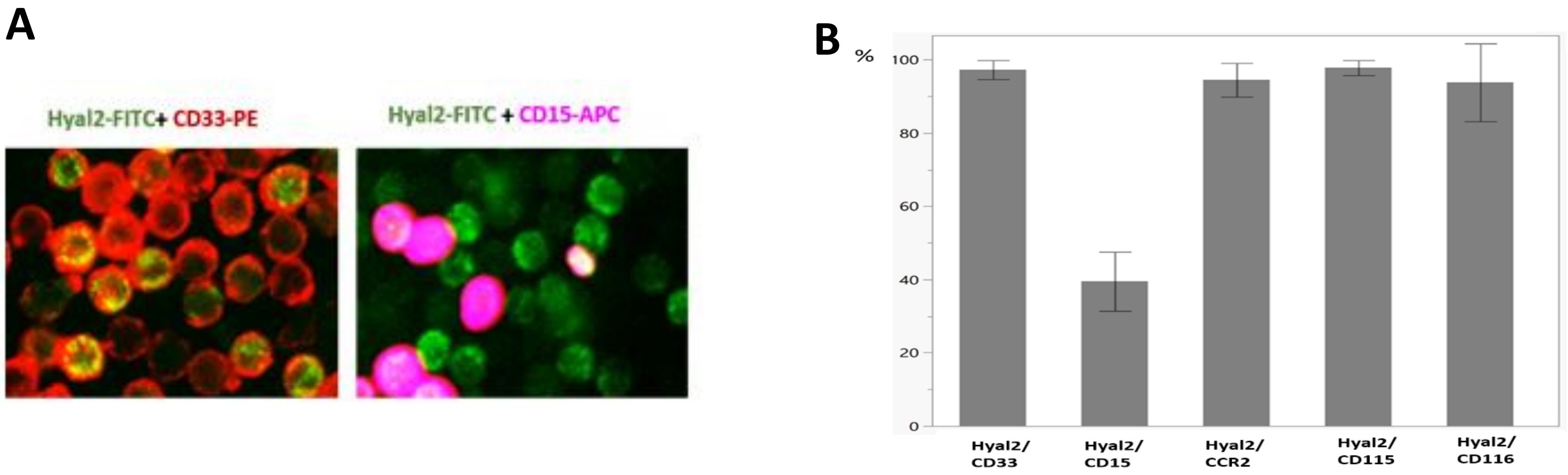


**Figure S7. Hyal2 myeloid cells express CD33, the marker of monocytic MDSCs. A**: CD11b myeloid cells were isolated from the freshly obtained peripheral blood of three bladder cancer patients, stained with anti-Hyal2-FITC and CD15-APC (right image), anti-Hyal2-FITC and CD33-PE (left image) and analyzed by IF microscopy. Representative images are shown. **B**: CD11b myeloid cells isolated from peripheral blood were stained with Hyal2-FITC antibody and with flurochrome-conjugated Abs against CD33, CD15, CCR2, CD115 and Cd116. Percent of double-positive cells was measured using microscope Lionheart FX and Gen 5 Prime v 3.08 software (Biotek Instruments).

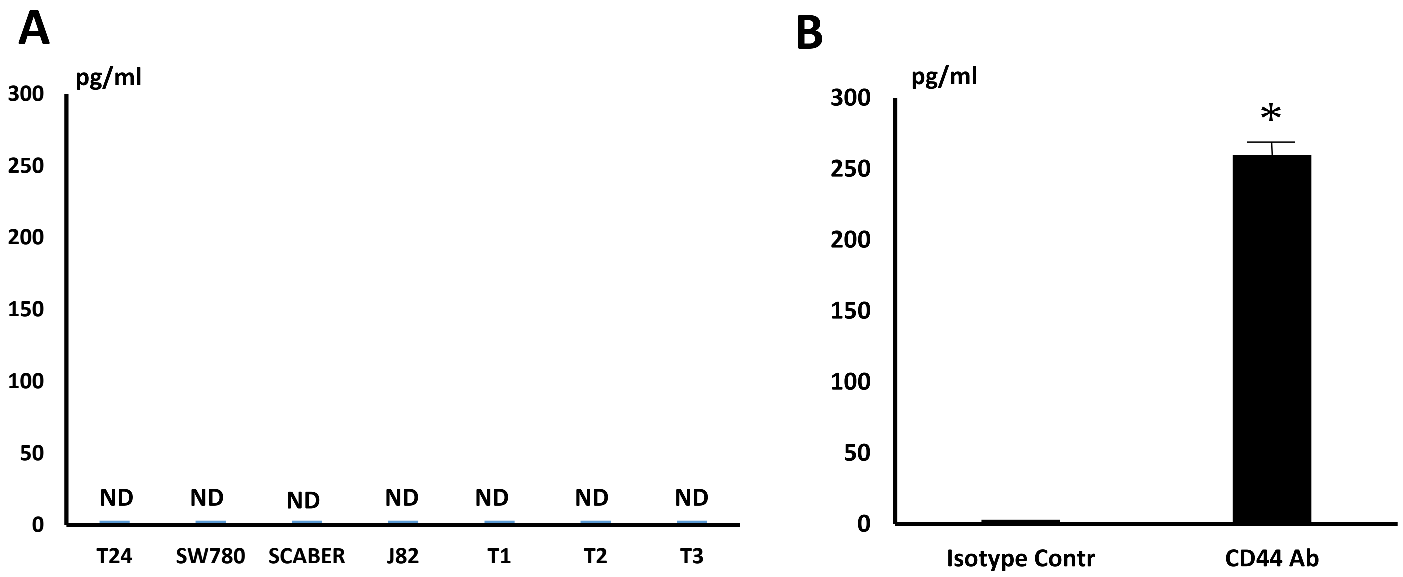


**Figure S8.** Stimulation of CD44 receptor With monoclonal antibody promotes IL-1beta production by blood-derived myeloid cells. **A:** Conditioned medium from human bladder cancer cell lines (T24, SW780, SCABER, J82) and from human bladder tumor tissue slice cultures (T1, T2 and T3) were analyzed for IL-1beta presence using ELISA. (**ND – Not Detected**)

**B:** $CD11b^{+}$cells were sorted from peripheral blood of bladder cancer patients using magnetic beads. Cells were plated in 24-well plates in complete culture medium and stimulated with 10 microm/ml of Isotype control Ab or CD44 Ab. Twenty four hours later supernatants were collected and stored at -80 C. IL-1beta levels measured by ELISA. Average means ± SD are shown, n=3; *, P<0.05.

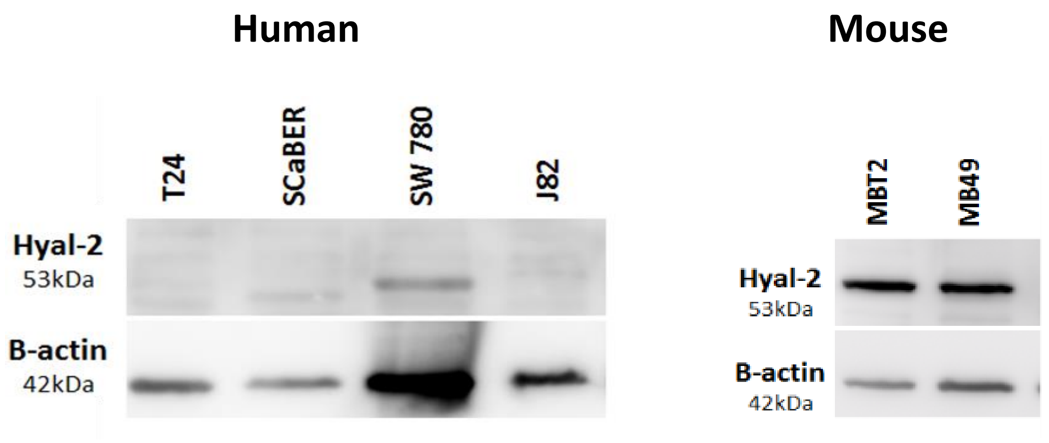


**Figure S9.Expression of Hyal2 in human and murine bladder tumor cell lines.** Cell lysates were prepared 4 human (T24, SCaBER, SW780, J82) and 2 mouse (MBT2 and MB49) bladder tumor cell lines. Cells were lysed in M-PER® mammalian protein extraction reagent (Thermo Scientific) containing protease and phosphatase inhibitors. Whole-cell lysates (30 µg/lane) were subjected to 10% SDS-PAGE, and blotted onto PVDF membranes. Results were visualized by chemiluminescence detection using a SuperSignal West Pico substrate (Thermo Scientific).
